# Global analysis of protein arginine methylation

**DOI:** 10.1101/2021.01.25.428036

**Authors:** Fangrong Zhang, Jakob Kerbl-Knapp, Maria J. Rodriguez Colman, Therese Macher, Nemanja Vujić, Sandra Fasching, Evelyne Jany-Luig, Melanie Korbelius, Katharina B. Kuentzel, Maximilian Mack, Alena Akhmetshina, Margret Paar, Beate Rinner, Gerd Hörl, Ernst Steyrer, Ulrich Stelzl, Boudewijn Burgering, Tobias Eisenberg, Brigitte Pertschy, Dagmar Kratky, Tobias Madl

**Affiliations:** Gottfried Schatz Research Center for Cell Signaling, Metabolism and Aging, Molecular Biology and Biochemistry, Medical University of Graz, 8010 Graz, Austria; Oncode Institute and Department of Molecular Cancer Research, Center for Molecular Medicine, University Medical Center Utrecht, 3584 CX, Utrecht, The Netherlands; BioTechMed-Graz, 8010 Graz, Austria; Institute of Pharmaceutical Sciences, University of Graz, 8010 Graz, Austria; Institute of Molecular Biosciences, NAWI Graz, University of Graz, 8010 Graz, Austria; Otto-Loewi Research Center, Physiological Chemistry, Medical University of Graz, 8010 Graz, Austria; Division of Biomedical Research, Medical University of Graz 8036 Graz, Austria; Field of Excellence BioHealth – University of Graz, Graz, Austria

**Keywords:** arginine methylation, NMR spectroscopy, protein arginine methyltransferases, mouse models, organoids, yeast, cancer, cell differentiation, ageing

## Abstract

Quantitative information about the levels and dynamics of post-translational modifications (PTMs) is critical for an understanding of cellular functions. Protein arginine methylation (ArgMet) is an important subclass of PTMs and is involved in a plethora of (patho)physiological processes. However, due to the lack of methods for global analysis of ArgMet, the link between ArgMet levels, dynamics and (patho)physiology remains largely unknown. We utilized the high sensitivity and robustness of Nuclear Magnetic Resonance (NMR) spectroscopy to develop a general method for the quantification of global protein ArgMet. Our NMR-based approach enables the detection of protein ArgMet in purified proteins, cells, organoids, and mouse tissues. We demonstrate that the process of ArgMet is a highly prevalent PTM and can be modulated by small-molecule inhibitors and metabolites and changes in cancer and during ageing. Thus, our approach enables to address a wide range of biological questions related to ArgMet in health and disease.

**Graphical Abstract:** 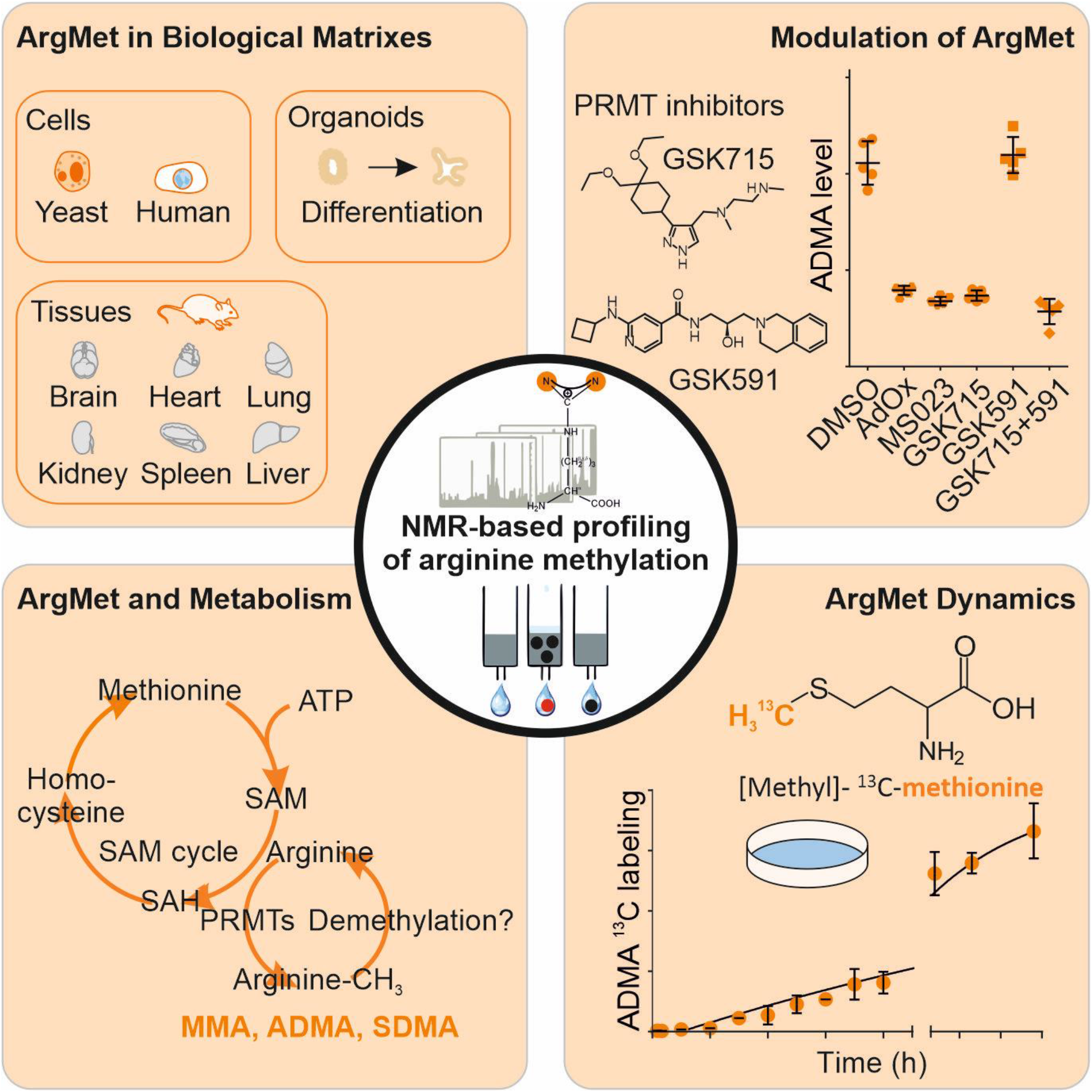

## Introduction

Arginine methylation (ArgMet) is a prevalent post-translational modification (PTM) evolutionary conserved from unicellular eukaryotes to humans. It regulates a plethora of fundamental biological processes, such as transcription, translation, RNA metabolism, signal transduction, DNA damage response, apoptosis, and liquid-liquid phase separation (LLPS) (Bachand, 2007; Bedford and Clarke, 2009; Bedford and Richard, 2005; Lee et al., 2005; Pahlich et al., 2006).

Three main types of methylated arginine residues are present in cells, including ω-*N*^*G*^-monomethylarginine (MMA), ω-*N*^*G*^, *N*^*G*^-asymmetric dimethylarginine (ADMA) and ω-*N*^*G*^-*N*’^*G*^-symmetric dimethylarginine (SDMA). Formation of MMA, SDMA and ADMA is catalysed by a broad spectrum of protein arginine methyltransferases (PRMTs). The number of PRMTs varies from unicellular eukaryotes to humans, with yeast having at least one or two main PRMTs (Hmt1/Rmt1 and Hsl7) and a family of nine PRMTs being present in mammals (Bachand, 2007; Bedford and Clarke, 2009). Depending on the type of methylated arginine they produce, PRMTs are categorized into four main classes (Bachand, 2007; Guccione and Richard, 2019). Type I PRMTs, including PRMT1, 2, 3, 4 (also called CARM1), 6, and 8 catalyse the formation of MMA/ADMA, whereas type II PRMTs, including PRMT5 and 9, catalyse the formation of MMA/SDMA (Figure S1A). Type III PRMTs such as PRMT7 catalyse the formation of MMA. In yeast, only the type IV PRMT Rmt2 has so far been reported (Chern et al., 2002) to methylate the delta (δ) nitrogen atom of arginine residues (Niewmierzycka and Clarke, 1999). Additional potential arginine methyltransferases have been identified, but remain to be biochemically validated (NDUFAF7, METTL23) (Guccione and Richard, 2019).

Most PRMTs methylate glycine- and arginine-rich, so-called arginine-glycine-glycine (RG/RGG), protein regions. More than 1000 human (in particular RNA-binding) proteins contain RG/RGG regions (Chong et al., 2018; Thandapani et al., 2013). In addition to the RG/RGG sites, PRMT4/CARM1 has been reported to methylate arginines within proline-, glycine-, and methionine-rich regions (Cheng et al., 2007). On a molecular level, methylation of these regions regulates nucleic acid binding, protein-protein-interactions, LLPS, and protein localization (Guccione and Richard, 2019).

PRMTs are ubiquitously expressed in human tissues, except PRMT8, which is primarily expressed in the brain, and regulate important cellular processes that affect cell growth, proliferation and differentiation. Embryonic loss in most of these PRMTs results in pre- and perinatal lethality. Dysregulation of PRMTs has been implicated in the pathogenesis of several diseases, including cardiovascular, metabolic, and neurodegenerative diseases, viral infections, and various types of cancer (Blanc and Richard, 2017). Since PRMTs tend to be upregulated in cancer malignancies (Jarrold and Davies, 2019; Yang and Bedford, 2013) they represent a promising target in cancer therapy and are currently being investigated in several clinical studies with PRMT inhibitors. Moreover, loss of PRMTs has been linked to cellular senescence and ageing in mice (Blanc and Richard, 2017).

Despite the biological significance of ArgMet, several key questions are still elusive: (i) The global levels of ArgMet are largely unknown. Pioneering studies indicated that ArgMet might be as abundant as phosphorylation, with around 0.5 - 2% of arginine residues being methylated in mammalian cells and tissues (Boffa et al., 1977; Esse et al., 2014; Matsuoka, 1972; Paik et al., 2007). However, specific concentrations of ArgMet in cells and tissues, including the coupling of ArgMet and metabolism, have so far not been comprehensively studied. The methyl group for protein ArgMet is provided by the universal methyl-donor S-adenosyl methionine (SAM), which is synthesized from methionine and ATP by SAM synthase. One carbon metabolism is required for recycling of the essential amino acid methionine (Locasale, 2013; Yang and Vousden, 2016). How metabolism regulates ArgMet needs to be determined. (ii) Dynamics and turnover of ArgMet, including the existence of an efficient arginine demethylase, are controversial issues and still largely unexplored (Guccione and Richard, 2019). (iii) Regulators of PRMTs (e.g. BTG1, TIS21/BTG2, or NR4A1) were proposed in the last years, but their impact on PRMT activity and, in turn, their contribution to global ArgMet concentrations remain enigmatic (Bedford and Richard, 2005; Yang and Bedford, 2013). (iv) Small-molecule inhibitors of PRMTs have been discovered, yet their influence on the extent of ArgMet and how ArgMet levels are affected *in vivo* is currently unknown.

Addressing these questions is challenging, in part due to the lack of robust methods for (absolute) quantification of global ArgMet values and dynamics in cells and tissues. Most of the current approaches use antibodies to detect methylated arginines. However, these antibodies are still only raised against very specific, short target sequences and fail to recognize or enrich larger numbers of arginine methylated proteins, limiting their use in quantification (Lee and Stallcup, 2009).

We therefore developed a general method for absolute, label-free quantification of (methylated) arginines in cells, organoids and tissues by using the high sensitivity and robustness of Nuclear Magnetic Resonance (NMR) spectroscopy. We demonstrate that ArgMet is a highly abundant PTM, whereas cellular dynamic changes of protein ArgMet occur at a slow rate. Our study provides a first strong methodological development for the quantification of ArgMet levels and their dynamic changes, that also conceptually advances our understanding of the importance of ArgMet in biology and medicine. Moreover, we offer new ways to study the modulation of protein ArgMet by inhibitors, metabolites and biological processes such as differentiation and ageing, enabling future studies from basic to translational research and drug discovery/development far beyond the current state of the art.

## Results

### NMR enables quantification of global protein arginine methylation

NMR spectroscopy enables robust quantification of metabolites in complex mixtures paired with simple and fast sample preparation, measurement and analysis (Stryeck et al., 2018). We built on previous chromatography-based approaches to analyse (methylated) arginines in protein hydrolysates (Dhar et al., 2013; Paik and Kim, 1967) and developed a novel NMR-based protocol for absolute quantification of protein ArgMet. A schematic representation of the workflow is shown in Figure 1A. Proteins were extracted from biological matrices, hydrolysed using hydrochloric acid, and delipidated. Basic/hydrophobic amino acids, including arginine and its derivatives, were purified by solid phase extraction (SPE) and analysed by NMR spectroscopy. NMR analysis of arginine, ADMA, MMA and SDMA standards revealed good separation of their ^1^H signals, both in 1D Car-Purcell-Meiboom-Grill (CPMG) as well as in 2D homo-nuclear J-resolved experiments (JRES) (Figure 1B, Figure 1C and Figure S1B). To minimise signal overlap with other metabolites present in biological materials, we used the JRES approach for all follow-up analyses. ^1^H-methyl signals of MMA and SDMA overlapped in ^1^H spectra when recorded in buffer, but could be resolved in d_6_-DMSO as solvent (Figure 1D).

**Figure 1.**
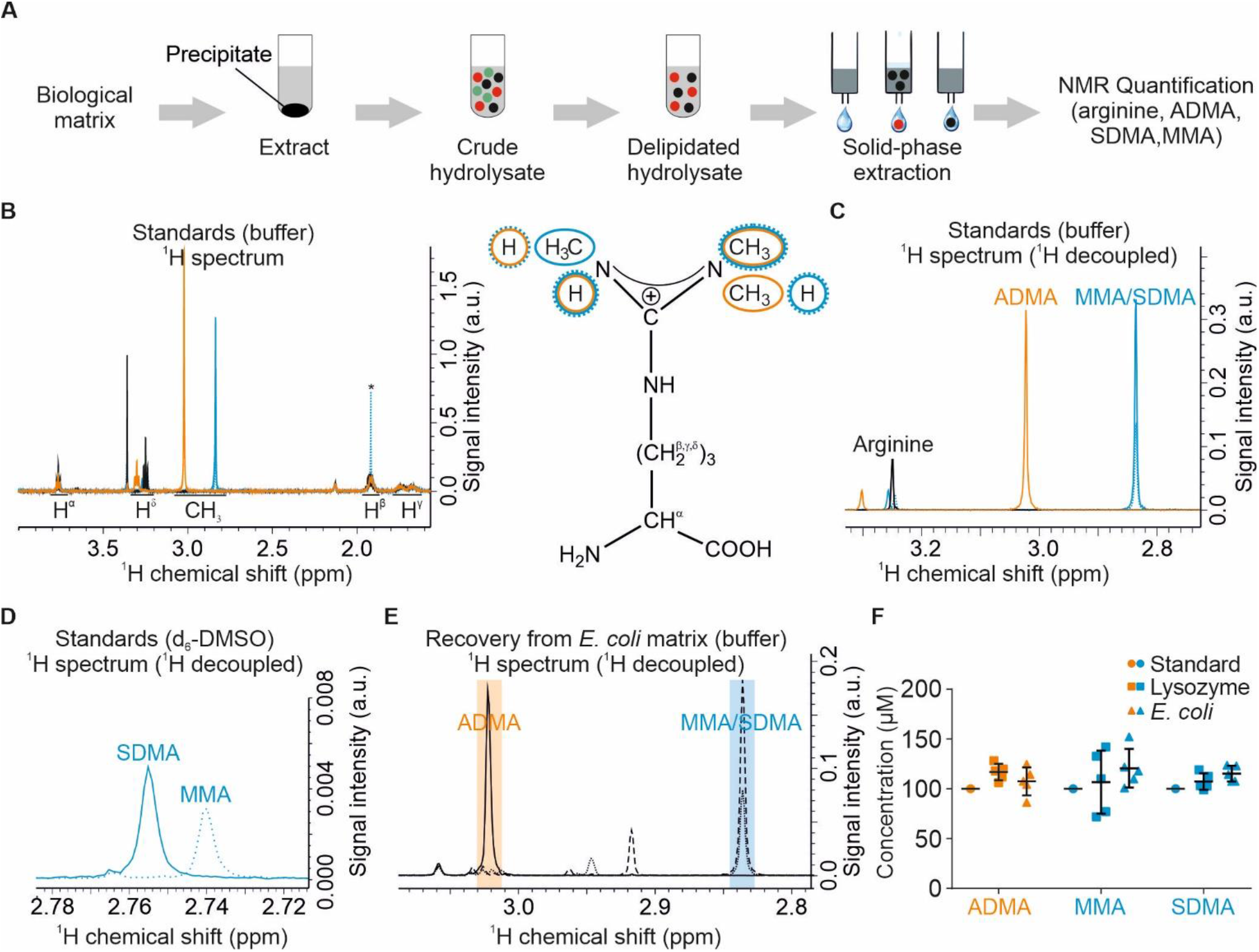
Absolute quantification of protein ArgMet by NMR. (A) Schematic workflow depicting steps for protein arginine and ArgMet quantification. Biological matrices are extracted with water/methanol. Protein precipitate containing protein arginine and ArgMet is hydrolysed, lipids are removed with chloroform, and solid phase extraction (SPE) is used to isolate positively charged amino acids, including (methylated) arginine(s). The eluate is analysed by NMR spectroscopy. (B) Overlay of ^1^H 1D-CPMG NMR spectra of 100 μM arginine (black), ADMA (orange), MMA (blue, dashed line) and SDMA (blue, solid line). Chemical shift ranges for characteristic ^1^H signals are shown in the spectra. Positions of the corresponding protons are labelled in the structure formula (ADMA - orange, MMA - blue, dashed line, SDMA-blue, solid line; an acetate impurity is labelled with an asterisk). (C) Overlay of ^1^H 1D projections of 2D J-resolved, virtually decoupled NMR spectra of the samples shown in (B). Characteristic regions of ADMA, MMA and SDMA methyl groups are indicated. (D) Overlay of ^1^H 1D projections of 2D J-resolved NMR spectra of 100 μM MMA and SDMA recorded in DMSO-d_6_ show the resolution of methyl resonances. (E) Overlay of representative recovery experiments of ^1^H 1D projections of 2D J-resolved NMR spectra recovery experiments from *E. coli* lysates spiked with ADMA (solid line), MMA (dotted line) or SDMA (dashed line), respectively. (F) Statistical analysis of ADMA, MMA, SDMA recovery from lysozyme (n=5; mean ± SD) (0.34 mM) and *E. coli* lysates (n=5, mean ± SD). Samples were spiked with 100 μM ADMA, SDMA, MMA and prepared according to the workflow shown in (A).

To validate the robustness of our workflow, we first evaluated stability and recovery of ADMA, MMA and SDMA signals in diverse biological matrices. All compounds were highly stable during hydrolysis and showed high recovery both from a protein matrix containing lysozyme and a methylation-free *Escherichia coli* cell matrix (Figure 1E, Figure 1F, Figures S1C-E). A quantitation limit for ADMA of 100 nM was determined (Figure S1F). Concentrations remained linear over a wide concentration range of 4 orders of magnitude up to the SPE column saturation limit of 3 mM, as shown for arginine (Figures S1G). In summary, our NMR approach offers a simple, rapid and highly reproducible workflow for arginine and ArgMet quantification. Compared to HPLC-based quantification, NMR is label-free, does not require chromatographic separation or standards for quantification. Moreover, it enables detection of yet unknown arginine derivatives and can be combined with isotope labelling.

### NMR-based protein ArgMet profiling *in vitro* and in cells

To identify the proportion of ArgMet in protein and cell samples of unknown methylation status, we determined levels of arginine, ADMA, MMA and SDMA in recombinant proteins, yeast cell cultures and mammalian cell lines. Levels of ADMA, MMA and SDMA are presented normalised to the total arginine content to allow a direct comparison of ArgMet concentrations between different biological matrices. Alternatively, and because NMR is completely quantitative, absolute concentrations can be displayed, normalised to either cell number, tissue mass or protein content.

Methylation by PRMTs occurs preferentially within RG/RGG-rich and proline-glycine-methionine-rich regions (Blanc and Richard, 2017). In mammals, PRMT1 is the most abundant methyl transferase and catalyses formation of both ADMA and MMA. As expected, NMR analysis of the methylation-free recombinant RG/RGG model proteins cold-inducible RNA-binding protein (CIRBP) and RNA-binding protein fused in sarcoma (FUS) revealed that ADMA and MMA are detectable in recombinant proteins after incubation with PRMT1 and the methyl-donor SAM (Figure 2A). Both model proteins are suitable as *in vitro* substrates for PRMT1, with 12% and 5% of all arginine residues being asymmetrically dimethylated in CIRBP and FUS, respectively. Interestingly, the levels of ADMA and MMA varied between CIRBP and FUS, with CIRBP lacking MMA/SDMA and FUS showing MMA/SDMA. These data indicate that PRMT1 has different sequence preferences for ADMA and MMA (Guo et al., 2014) and that ArgMet-NMR is well-applicable to study levels and kinetics of ArgMet in purified protein substrates.

**Figure 2.**
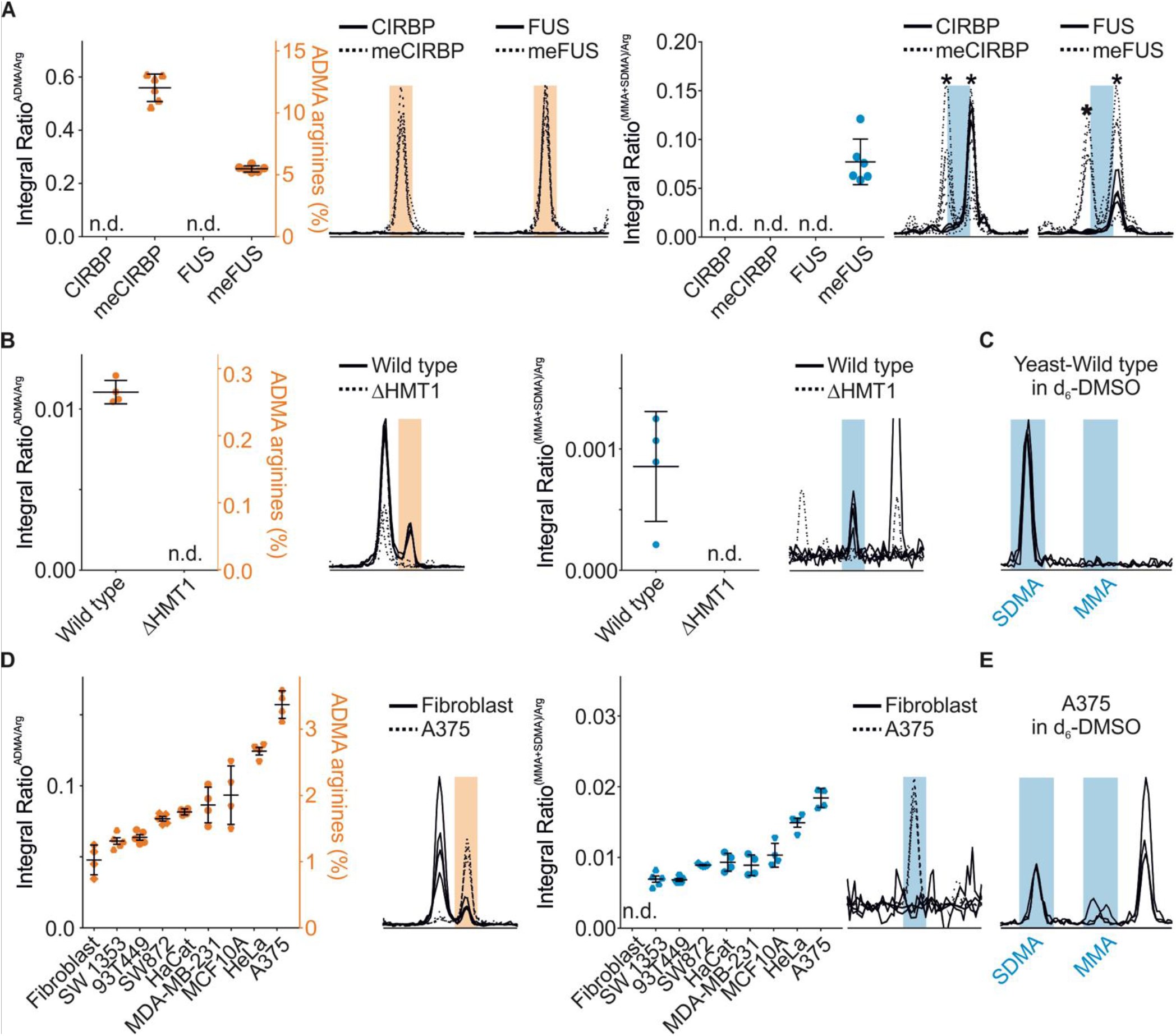
Characterisation of ArgMet in purified proteins, yeast and mammalian cell lysates. (A) ArgMet quantification of recombinant CIRBP and FUS peptides without methylation or *in vitro* methylated by recombinant PRMT1, respectively (n=6; mean ± SD; n.d. - not detectable; Tris buffer impurities are labelled with asterisks). (B) Protein ArgMet quantification of yeast lysates (n=4; mean ± SD; n.d. - not detectable). (C) Spectral overlays of characteristic MMA and SDMA NMR methyl signals in d_6_-DMSO show that MMA and SDMA methyl resonances can be resolved (n=3). (D) Protein ArgMet quantification of human cell lysates (n=4-5; mean ± SD; n.d. - not detectable). ADMA levels in relation to the total amount of arginine are indicated. Spectral overlays of characteristic ADMA and MMA/SDMA NMR methyl signals are shown (n=4). (E) Spectral overlays of characteristic MMA and SDMA NMR methyl signals in d_6_-DMSO show that MMA and SDMA methyl resonances can be resolved (n=3). In A, B and D, the ADMA concentrations are indicated relative to the total amount of arginine. Spectral overlays of characteristic ADMA (orange) and MMA/SDMA (blue) NMR methyl signals are shown (n=4-5).

First HPLC-based studies estimated 0.5 - 2% of arginine residues to be methylated in mammalian cells and tissues (Boffa et al., 1977; Esse et al., 2014; Matsuoka, 1972; Paik et al., 2007). In yeast, however, ArgMet has not been well-studied, although four PRMTs have been described in *Saccharomyces cerevisiae* (Hmt1/Rmt1, Rmt2, Hsl7 and Sfm1). Of these, Hmt1/Rmt1 has been identified as PRMT1 homologue (Bachand, 2007). Analysis of wild-type and *HMT1/RMT1* knock-out yeast strains showed that on average more than 0.25% of all arginines are methylated in *S. cerevisiae* (Figure 2B). SDMA was detectable in wild-type yeast, albeit at low levels (~10% of ADMA), whereas MMA was undetectable (Figure 2C). Deletion of *HMT1/RMT1* essentially abolished ADMA and SDMA levels, suggesting that none of the other PRMTs contributed significantly to the global ArgMet levels. In line with these results, only very few substrates of Rmt2, Hsl7 and Sfm1 have been reported so far (Chern et al., 2002; Sayegh and Clarke, 2008; Young et al., 2012). Nevertheless, we cannot exclude indirect effects, e.g. a possible downregulation of other PRMTs by the loss of Hmt1/Rmt1.

Our approach offers an excellent opportunity to characterise ArgMet in a variety of commonly used human cell lines. ArgMet-NMR analysis of nine human cells lines showed that ADMA and MMA/SDMA concentrations differ significantly, with primary fibroblasts showing the lowest and A375 malignant melanoma cells showing the highest levels of both ADMA and MMA/SDMA, respectively (Figure 2D). In all cell lines tested, ADMA was the predominant ArgMet species with more than 3% of all arginine residues being methylated in A375 cells. SDMA/MMA levels were significantly lower (Figures 2D and E). This finding is in line with previous studies estimating MMA and SDMA at levels of 20-50% of ADMA (Bedford and Clarke, 2009), although the MMA/SDMA values detected by NMR are consistently lower (~10% of ADMA). The significantly increased concentrations of ArgMet in A375 cells compared to all other cell lines is in agreement with a recent study showing overexpression of PRMT1 in these cells (Li et al., 2016). In contrast, HPLC-based methods detected 0.8% of all arginine residues in A375 cells being asymmetrically dimethylated, which is lower than the 3.4% of ADMA we found (Bulau et al., 2006). The increase of ADMA in all investigated cell lines is correlated with a concomitant increase in MMA and SDMA, indicating that the corresponding enzymes are co-regulated (Figure S2). Generally, non-cancer cell lines such as primary fibroblasts show a tendency to lower concentrations of ArgMet compared to cancer cell lines such as HeLa, A375 or MDA-MB-231. Although HaCaT cells, an immortalized human keratinocyte line, exhibit also higher ArgMet levels, the values are still lower than in cancer cell lines. In summary, our approach provides a direct read-out of protein ArgMet in cell lines.

### NMR reveals modulation and dynamics of protein ArgMet

Although it has taken 50 years to acknowledge the significance of PRMTs in cancer, the pace at which major discoveries have been made in recent years is phenomenal. Disruption of ADMA modification at key substrates decreases the metastatic and proliferative ability of cancer cells (Li et al., 2016), suggesting that PRMT inhibitors may be an effective strategy to combat different types of cancer. Several PRMT inhibitors have entered or are on the verge of entering the clinic, but how they alter global protein ArgMet levels remains to be uncovered.

Since our method provides a first direct read-out of ArgMet modulation by PRMT inhibitors, we characterized ArgMet concentrations under distinct conditions of PRMT inhibition (Figures 3A and B). We first tested the impact of the commonly used general ArgMet inhibitor adenosine dialdehyde (AdOx) and the type I PRMT inhibitor MS023 (Afman et al., 2005; Chan-Penebre et al., 2015; Guccione and Richard, 2019). In line with our hypothesis, AdOx unselectively, though incompletely, reduced any kind of protein ArgMet significantly by ~60% (p<0.0001). As expected for a selective type I PRMT inhibitor, MS023 inhibited mostly ADMA, but not SDMA formation. PRMT5, the major enzyme catalysing the formation of SDMA, has been implicated in cancer biology, and controls expression of both tumour-suppressive and tumour-promoting genes (Guccione and Richard, 2019). Inhibition of PRMT5 by the small-molecule compounds GSK3203591 or GSK3326595 has been reported to act anti-proliferatively on mantle cell lymphoma, both *in vivo* and *in vitro* (Chan-Penebre et al., 2015; Gerhart et al., 2018). Moreover, GSK3368715, a reversible type I PRMT inhibitor, exhibited anti-tumour effects in human cancer models and is currently in phase I clinical trials (Guccione and Richard, 2019). GSK3203591 and GSK3368715 have been reported to synergistically inhibit tumour growth *in vivo*, possibly through a tumour-specific accumulation of 2-methylthioadenosine, an endogenous inhibitor of PRMT5, which correlates with sensitivity to GSK3368715 in cell lines (Fedoriw et al., 2019). In agreement, GSK3368715 inhibited formation of ADMA, but not SDMA formation (Figure S3A), whereas GSK3203591 inhibited generation of MMA/SDMA but not of ADMA. Compared to all other conditions tested, a combination of GSK3203591 and GSK3368715 showed the strongest inhibition of any type of ArgMet in HeLa cells. Interestingly, inhibition of type I PRMTs by MS023 or GSK3368715 doubled the levels of SDMA/MMA, suggesting that, on a global scale, several type I PRMT targets become symmetrically instead of asymmetrically dimethylated. Accordingly, recent Western blotting experiments resulted in increased MMA/SDMA levels after treatment with PRMT1 inhibitors (Eram et al., 2016; Fedoriw et al., 2019). Comparable results in other cell lines demonstrated that the mechanisms of ArgMet inhibition are independent of the cell line (Figures S3B and C).

**Figure 3.**
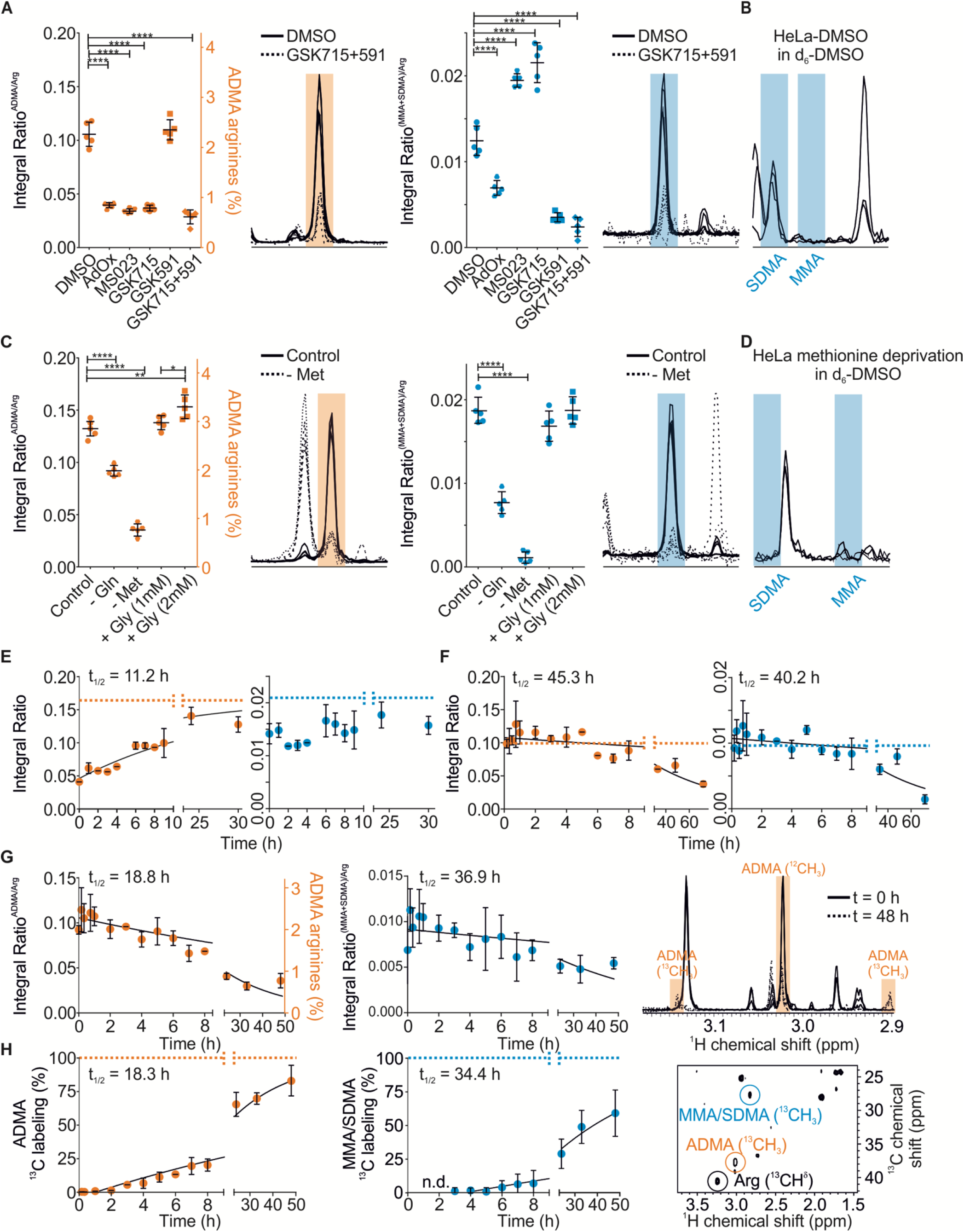
NMR enables quantification of protein ArgMet modulation and dynamics. (A) Protein ArgMet quantification of HeLa cells treated for 3 days with either DMSO, 40 μM adenosine dialdehyde (AdOx), 10 μM MS023, 2 μM GSK3368715 (GSK715), 1 μM GSK3203591 (GSK591), or a combination of 2 μM GSK715 and 1 μM GSK591 (n=5; mean ± SD). ADMA levels in relation to the total amount of arginine are indicated. Spectral overlays of characteristic ADMA (orange) and MMA/SDMA (blue) NMR methyl signals are shown (n=5). (B) Spectral overlays of characteristic MMA and SDMA NMR methyl signals in d_6_-DMSO show that MMA and SDMA methyl resonances can be resolved (n=3). (C) Protein ArgMet quantification of HeLa cells cultured with or without 4 mM glutamine (Gln), 0.2 mM methionine (Met) or glycine (1 mM and 2 mM Gly) (n=5; mean ± SD). ADMA levels in relation to the total amount of arginine are shown. Spectral overlays of characteristic ADMA (orange) and MMA/SDMA (blue) NMR methyl signals are presented (n=5). (D) Spectral overlays of characteristic MMA and SDMA NMR methyl signals in d_6_-DMSO show that SDMA levels strongly decrease upon methionine deprivation (n=3). (E) Changes of ArgMet levels after removal of AdOx. Prior to removal of AdOx, HeLa cells were treated with AdOx for 3 days to reduce ArgMet. Integral ratios of ADMA/arginine and (SDMA+MMA)/arginine are plotted as mean ± standard error (n=3) for each time point. To estimate the half-life of ArgMet recovery (t_1/2_) the data were fitted using a single exponential recovery function (95% CI: 7.5-19.9 h). Dotted lines indicate the level of methylation in the absence of AdOx. (F) Changes of ArgMet levels after methionine removal. Integral ratios of ADMA/arginine and (SDMA+MMA)/arginine are plotted as mean ± standard error (n=3) for each time point. To estimate the half-life of ArgMet decay (t_1/2_) the data were fitted using a single exponential decay function (ADMA/arginine - 95% CI: 34.0-62.7 h; (SDMA+MMA)/arginine - 95% CI: 27.8-61.1 h). Dotted lines indicate the level of methylation in the presence of methionine. (G) Dynamics of *de novo* ArgMet via ^13^C labelling are shown as decay of the ^12^C-methyl NMR signals upon exchange of media containing ^13^C-methyl-labelled methionine. Integral ratios of ADMA/arginine and (SDMA+MMA)/arginine are plotted as mean ± standard error (n=3) for each time point. To estimate the half-life of ArgMet ^12^C-methyl signal decay (t_1/2_), the data were fitted using a single exponential decay function (ADMA/arginine, 95% CI: 14.4-24.7; (SDMA+MMA)/arginine, 95% CI:21.9-73.8). Change of ADMA ^12^C/^13^C-methyl signals at the beginning and after 48 h of cultivation in presence of ^13^C-methyl-labelled methionine (^1^H 1D projections of 2D J-resolved NMR spectra). (H) Dynamics of *de novo* ArgMet via ^13^C labelling are shown as increase of the ^13^C-methyl NMR signals detected in ^1^H,^13^C HSQC NMR spectra upon exchange of media containing ^13^C-methyl-labelled methionine. Fractions of ^13^C-labeling are plotted as mean ± standard error (n=3) for each time point. To estimate the half-life of ArgMet ^13^C-methyl labelling (t_1/2_), the data were fitted using a one phase association function (ADMA/arginine, 95% CI: 16.4-21.2; (SDMA+MMA)/arginine, 95% CI:27.4-43.5). Arginine, ADMA, MMA, and SDMA ^1^H,^13^C NMR signals are labelled in a representative ^1^H,^13^C HSQC NMR spectrum.

One of the main goals of current ArgMet research is to further refine our mechanistic understanding of ArgMet and how this process is coupled with metabolism. PRMTs add methyl groups to arginine residues using the universal methyl-donor SAM, which is recycled through one-carbon metabolism (Ducker and Rabinowitz, 2017; Yang and Vousden, 2016). Methionine is a key substrate for SAM production. Beyond the single-carbon metabolic pathway, additional metabolites can alter global cellular ArgMet by modulating the SAM levels and recycling. Indeed, considering that methionine is an essential amino acid and its recycling can therefore only partly contribute to the methionine pool required for SAM generation, deprivation of methionine strongly reduced the concentrations of ADMA and MMA/SDMA in HeLa cells (Figures 3C and D). Production of SAM requires ATP, and its recycling via S-adenosyl-homocysteine depends on supply of the single-carbon building block from serine (Yang and Vousden, 2016). In cancer cells, glutamine can provide both the single-carbon building block through gluconeogenesis and energy through the tricarboxylic acid cycle, respectively. Thus, we tested in HeLa cells if depletion of glutamine reduced the overall levels of protein ArgMet. In line with our hypothesis, concentrations of ADMA and SDMA/MMA were reduced, although not as profoundly as in the case of methionine withdrawal. Glycine supplementation has been proposed to mimic the effects of methionine deprivation through inhibition of the serine-to-glycine conversion that otherwise provides the single-carbon building block for SAM recycling (Partridge et al., 2020). In contrast to these studies, we observed even an increase in ADMA when cells were incubated with 2 mM glycine (Figure 3C). Taken together, ArgMet-NMR provides a novel toolbox for future studies of protein ArgMet regulation by inhibitors and metabolites.

Dynamics of arginine methylation and demethylation is one of the yet unsolved and controversial questions in the field, in particular it is still unclear, whether an efficient arginine demethylase exists (Guccione and Richard, 2019). We therefore first addressed the dynamics of re-methylation in a low-ArgMet background. We treated cells with medium supplemented with AdOx to reduce ArgMet, then exchanged the medium to AdOx-free medium and collected cells at different time points. We found that levels of ArgMet recovered slowly after AdOx removal, with ADMA having a half-life of > 11 hours (Figure 3E) As this process might have been affected by the levels of AdOx decreasing slowly inside the cell, we further validated the changes in ArgMet concentrations using methionine deprivation. Under these conditions, levels of protein ArgMet decreased considerably (~60%), with half-lives of 45 and 40 hours for ADMA and MMA/SDMA, respectively (Figure 3F). Although these alterations are strongly coupled to the dynamics of the cellular pool of methionine, our results indicate that de-methylation of methylated arginine residues is a slow process and that the available levels of methionine are insufficient to maintain the methylation levels over a longer period of time.

To monitor the dynamics of ArgMet in the absence of any interferences due to the manipulation of metabolic pathways, we combined ArgMet-NMR with stable isotope tracing using ^13^C-methyl-labelled methionine. With methionine being an essential substrate for SAM production, we next examined whether the methyl group crucial for ArgMet is donated by methionine and investigated the dynamics of the associated methylation reaction. To track and quantify *de novo* ArgMet, we pulsed HeLa cells in media with ^13^C-methyl-labelled methionine and chased its appearance by the decay of the ^12^C-methyl NMR signals upon exchange with media containing ^13^C-methyl labelled methionine (Figures 3G and H). Coupled with the decrease of ^12^C protein ArgMet, ‘newly’ synthesized and ^13^C isotopically labelled protein ArgMet appears (Figure 3H). Fitted half-lives of de-methylation (^12^C-decay) and *de novo* methylation (^13^C-increase) were in excellent agreement and approximately 18-19 and 34-37 hours for ADMA and MMA/SDMA, respectively. In line with the AdOx removal and methionine deprivation changes, these data indicate that the overall dynamics of arginine demethylation are slow.

### NMR provides first insights into dynamics of ArgMet in organoids and tissues

Increasing evidence suggests that ArgMet is required to maintain cells in a proliferative state and plays a key role in the homeostasis of stem cell pools (Blanc and Richard, 2017). In addition, the role of PRMTs has been associated with cell growth, differentiation, apoptosis and ageing (Blanc and Richard, 2017; Guccione and Richard, 2019; Yang and Bedford, 2013). For example, depletion and exhaustion of muscle and hematopoietic stem cells in adulthood was linked to loss of ArgMet (Blanc et al., 2016; Liu et al., 2015). In addition, PRMTs play important regulatory roles in the differentiation of myeloid cells (Balint et al., 2005). To study the relationship of ArgMet and *in vitro* differentiation in a controlled manner, we generated cell type-enriched mouse small intestinal organoid cultures. We grew the organoids in complete ENR medium (EGF, Noggin, R-Spondin) as reference. In ENR medium, organoids contain stem cells, enterocytes and Paneth cells (roughly 15%, 80%, 5%, respectively). Stem cells, enterocytes and Paneth cells were enriched using media supplemented with Wnt-CM (conditioned medium)/valproic acid (VPA; stem cells enriched), removal of R-Spondin (EN; enterocytes enriched), or supplementation of Wnt-CM/N-[N-(3,5-difluorophenacetyl)-L-alanyl*]-S-phenylglycine t-butyl ester) (DAPT; Paneth cells enriched), respectively. Our data show alterations of ADMA and MMA/SDMA dependent on organoid composition (Figure 4A). In line with a high expression of PRMTs in stem cells found in single-cell mRNA sequencing of mouse small intestine (Haber et al., 2017; Ludikhuize et al., 2020; Uhlen et al., 2015)(Figure S4, http://www.proteinatlas.org/), reference organoids (ENR) show higher ADMA and MMA/SDMA. Enrichment of Paneth cells in organoids (DAPT) results in a strong decrease in overall ArgMet, in line with a low expression of PRMTs in Paneth cells (Figure S4). However, it remains to be investigated whether ArgMet is a cause or consequence of differentiation and to elucidate the key regulatory and metabolic mechanisms modulating ArgMet during differentiation.

**Figure 4.**
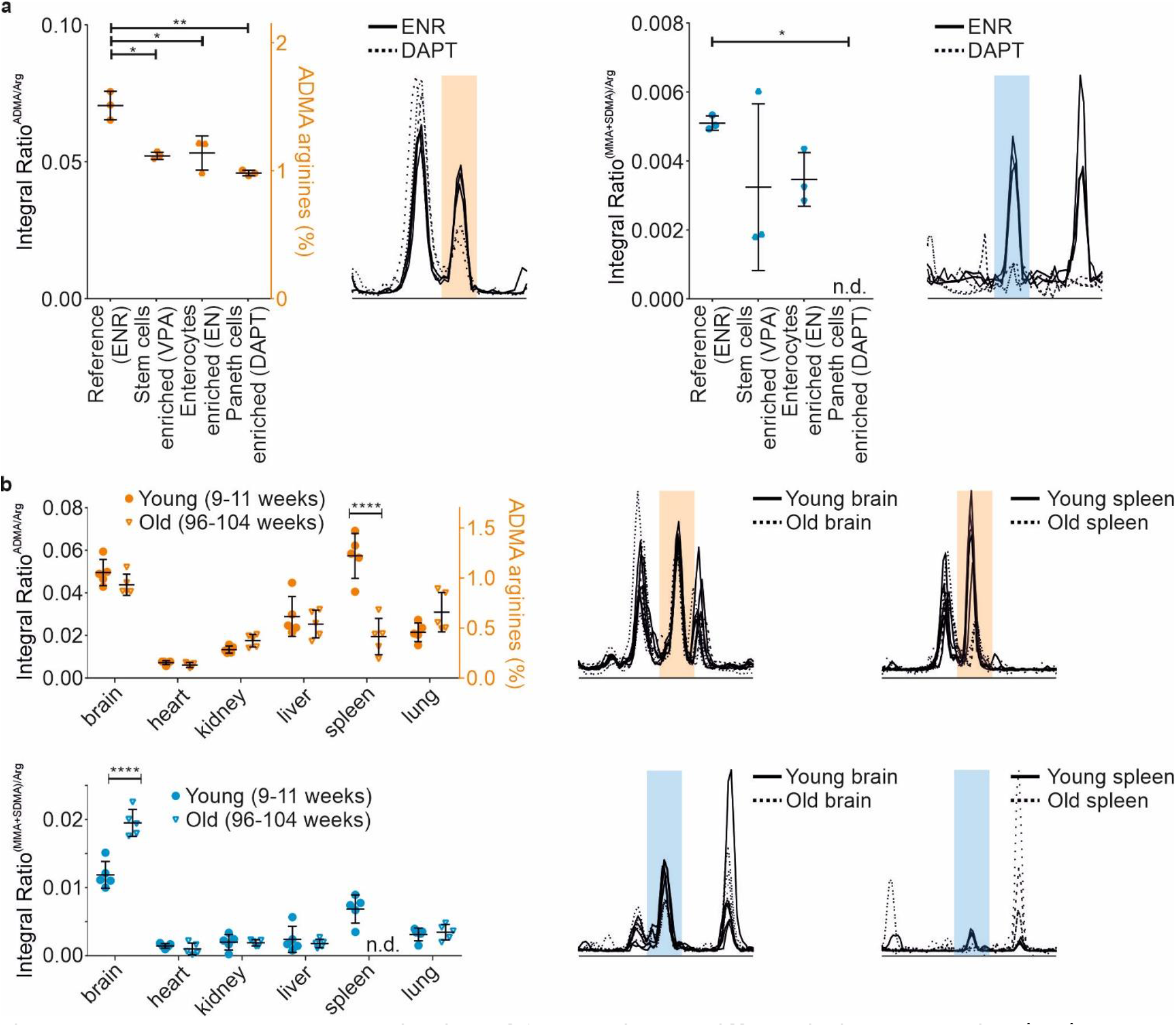
NMR enables characterization of ArgMet in cell differentiation and ageing *in vivo*. (A) Protein ArgMet quantification of human colon organoids cultured with EGF/Noggin/R-spondin1 (ENR), EGF/Noggin (EN), ENR plus valproic acid (VPA), or ENR plus Notch pathway inhibitor DAPT (N-[N-(3, 5-difluorophenacetyl)-L-alanyl*]-S-phenylglycine t-butyl ester) (n = 3; mean ± SD). ADMA levels are presented with respect to total amounts of arginine. Spectral overlays of characteristic ADMA (orange) and MMA/SDMA (blue) NMR methyl signals are shown (n=3). (B) Quantification of protein ArgMet in mouse tissues collected from young (9-11 weeks, dots) and old mice (96-104 weeks, triangles) (n=5; mean ± SD). ADMA levels are presented with respect to total amounts of arginine. Spectral overlays of characteristic ADMA (orange) and MMA/SDMA (blue) NMR methyl signals are shown (n=3).

Studying the global levels of ArgMet *in vivo* is the ultimate goal to reveal the mechanistic links between ArgMet and (patho)physiology. To demonstrate the feasibility of our approach for *in vivo* studies, we characterised ArgMet levels in commonly studied mouse tissues (brain, heart, kidney, liver, spleen and lung) in two groups of female wild-type mice (mixed background of 129/J and C57BL/6J) at young (9-11 weeks) and old (96-104 weeks) age. Strikingly, we observed varying levels of ADMA and MMA/SDMA among tissues, with highest levels of ArgMet in brain and spleen of young mice (Figure 4B). Recent studies revealed high expression of PRMT1 in the rat spleen and a high expression of PRMT5 in the rat brain (Hong et al., 2012), substantiating our findings of high ADMA in the spleen and high MMA/SDMA in the brain. Moreover, PRMT1 and PRMT8 expressions were elevated in mouse brain compared to liver (Wang et al., 2017). With the exception of brain and spleen, we observed no significant ageing-related changes in ArgMet levels in any other tissues. Expression and catalytic activity of PRMT1, PRMT4, PRMT5, and PRMT6 have been reported to be reduced in replicatively senescent cells relative to young cells (Lim et al., 2010; Lim et al., 2008). In addition, senescent cells accumulate in tissues with age along with a decline in immune function (Kuilman et al., 2008). Although the underlying molecular mechanisms remain elusive, one might speculate that the changes in the aged spleen are caused by the accumulation of senescent cells.

## Discussion

Protein ArgMet modulates the physicochemical properties of proteins and thus plays a major role in a multitude of regulatory pathways, including gene regulation, signal transduction, regulation of apoptosis and DNA repair (Blanc and Richard, 2017; Guccione and Richard, 2019; Yang and Bedford, 2013). Although previous studies exist in this field, the lack of a reliable quantification of ArgMet is a restricting factor in elucidating the relevance of ArgMet in physiological and pathological processes. We have developed a simple, fast and robust protocol for NMR-based quantification of protein ArgMet levels and dynamics in purified proteins, cells, organoids and tissues. Our study reveals that NMR spectroscopy provides a sensitive read-out for detection and quantification of MMA, ADMA and SDMA in all matrices tested. We show that ArgMet-NMR enables detection of methylation patterns in purified proteins incubated with PRMT1. Methylation by PRMTs in the human proteome occurs preferentially (but not exclusively) within glycine-arginine- and proline-glycine-methionine-rich regions (Cheng et al., 2007; Thandapani et al., 2013; Woodsmith et al., 2018), but specific consensus sequences targeted by most of the human PRMTs remain to be identified. Our approach provides a toolbox for fast and label-free screening for PRMT selectivity in purified proteins/peptides.

Although some of human PRMTs are well-studied, it is yet unknown for a plethora of PRMTs from other organisms whether they exhibit any enzymatic activity (Fulton et al., 2019). For example, the main yeast methyltransferase is Hmt1, the presumable ortholog of human PRMT1. In addition, *in silico* studies have predicted 33 additional putative methyltransferases in *S. cerevisiae,* and it is likely that beside nucleic acid methyltransferases, and protein methyltransferases specific to other amino acids, also arginine methyltransferases are among them (Low and Wilkins, 2012). We found that *S. cerevisiae* produces ADMA and SDMA, but no MMA. This is in contrast to a report which identified low levels of MMA, but no SDMA in *S. cerevisiae* (Hsieh et al., 2007). According to our data, low concentrations of SDMA need a highly sensitive method like ArgMet-NMR to detect SDMA in yeast. Strikingly, deletion of *HMT1* led to a complete loss of ADMA and MMA, suggesting that the contribution of any other methyltransferase to global levels of ArgMet are negligible, at least in this yeast strain. However, we cannot exclude the possibility that the other putative methyltransferases methylate only a small subset of targets, resulting in low global ArgMet levels. Our proof-of-principle analysis in human cell lines identified a large fraction of arginine residues in a methylated state, ranging from 1 to 3.4%. In all cell lines tested, ADMA constituted the predominant methylated arginine species, followed by SDMA with about 10% and MMA with about 1% of ADMA. These ADMA levels are in agreement with the findings that PRMT1 is the predominant and most active PRMT present in mammalian cells (Tang et al., 2000). Screening the PhosphoSitePlus(R) database of PTMs for ArgMet revealed that for 1.7% of all arginines in human proteins methylation (ADMA, SDMA or MMA) has been reported. Given that we identified between a methylation status of 1 and 3.4% of arginines to be methylated indicates that most of the proteins for which ArgMet has been reported are entirely methylated. Note that this estimation assumes that all proteins are present at comparable levels inside the cell (Hornbeck et al., 2015). Methylarginines are predominantly found in intrinsically disordered protein regions, e.g. RG/RGG regions, which are intimately connected to LLPS (Chong et al., 2018; Guccione and Richard, 2019; Woodsmith et al., 2018). A large proportion of the proteins implicated in LLPS are known targets for ArgMet, and therefore LLPS could be regulated by their ArgMet (Chong et al., 2018; Guccione and Richard, 2019; Lorton and Shechter, 2019). Thus, it is conceivable that the global ArgMet levels regulate LLPS on a global scale *in vivo* by regulating fluidity and dynamics of membrane-less organelles containing e.g. RG/RGG proteins.

By comparing the methylation levels in cell lines, ArgMet were up to 3-fold higher in immortalized and cancer cells compared to primary cells. Strikingly, cells isolated from human metastases contained the highest levels of protein ArgMet. The increased ArgMet levels found in cancer cells are in line with overexpression of PRMT1 in human melanoma, breast and prostate cancer (Bedford, 2007; Hamamoto and Nakamura, 2016). In addition, PRMT5 expression and activity seem to be important in tumorigenesis and are markers of poor clinical outcome (Stopa et al., 2015). Based on the observation that increased PRMT expression is associated with tumour growth, inhibitors of protein arginine methyltransferases have been developed and showed promising results in clinical studies (https://clinicaltrials.gov). Our study demonstrates that ArgMet-NMR provides a precise and specific read-out for modulation of ArgMet levels in cells treated with distinct (specific) PRMT inhibitors. This suggests that ArgMet-NMR might be a valuable tool for ArgMet-based drug discovery, drug validation and patient stratification in the future.

By examining the modulation of ArgMet levels upon metabolite deprivation in cells, we detected a tight metabolic regulation of ArgMet levels by methionine, glutamine and glycine. Methionine is required for protein synthesis and its adenylation produces SAM, which serves in turn as a methyl donor for methylation reactions (Locasale, 2013; Yang and Vousden, 2016). Accordingly, we demonstrated that methionine deprivation had a strong impact on protein ArgMet by reducing ADMA, MMA and SDMA by more than 61%. Given that methionine is an essential amino acid whose levels are dictated by dietary factors (Mentch and Locasale, 2016), it is conceivable that nutrition and fasting could, in addition to protein synthesis, additionally affect protein ArgMet *in vivo*. Moreover, glutamine deprivation in HeLa cells culture reduced protein ArgMet by more than 30%, corroborating the observation that glutamine is a key energy source in cancer cells and can provide the single-carbon building block for SAM recycling through gluconeogenesis (Curi et al., 2005). Glutamine plays a pleiotropic role in cellular function and its consumption is elevated in proliferating cells not only due to increased DNA production (Counihan et al., 2018; Vander Heiden and DeBerardinis, 2017), but also for maintaining high ArgMet levels. Notably, ArgMet requires an energy demand of 12 molecules of ATP per methylation event (Gary and Clarke, 1998). Thus, reduced energy supply by glutamine deprivation could be the major factor of the observed reduction in protein ArgMet. Glycine is an interesting metabolite due to its role in SAM recycling and methionine clearance. On the one hand, glycine can act as methyl group acceptor, leading to the formation of sarcosine (N-methylglycine) and S-adenosylhomocysteine. On the other hand, glycine is converted when the single-carbon block is transferred to tetrahydrofolate, which in turn is used to recycle SAM (Ducker and Rabinowitz, 2017; Luka et al., 2009; Yang and Vousden, 2016). Excess glycine has been proposed to reduce methionine levels and to mimic methionine deprivation (Partridge et al., 2020).

According to our findings but in contrast to previous studies, glycine supplementation failed to reduce global levels of protein ArgMet. The fact that glycine supplementation did not alter methionine levels in adult worms (Liu et al., 2019) suggests that under physiological conditions glycine supplementation is not generally applicable to mimic methionine deprivation in cancer cells. Our approach is expected to substantiate specific aspects of protein methylation research in the future. Moreover, it will be interesting to reveal whether lifespan extension via methionine restriction is mediated via modulation in ArgMet (Bárcena et al., 2018; Grandison et al., 2009; Lee et al., 2016).

Dynamics of cellular protein ArgMet, the process of methylation and the process of ‘demethylation’, can also be easily examined by our novel methodology. The existence of an efficient arginine demethylase has not yet been proven and is a long-disputed question in this field (Low et al., 2016). Our results obtained using different setups of re-methylation after treatment with the general methylation inhibitor AdOx and ‘demethylation’ upon methionine deprivation show that global arginine (de)methylation is a slowly developing process in a cellular context. We further substantiated these findings by combining ArgMet-NMR with stable isotope tracing using ^13^C-methyl-labelled methionine. A decrease of the NMR signal characteristic for unlabelled methylated arginine residues in combination with an increase of the NMR signal characteristic for ^13^C-methylated arginines indicated that both *de novo* ArgMet and ‘demethylation’ are slow processes, especially in comparison to phosphorylation and de-phosphorylation. For example, global phosphorylation of the epidermal growth factor receptor occurs within 2–3 hours with a half-life of approximately 30 min, whereas its intracellular domain is dephosphorylated considerably faster (t_1/2_ = 15 s) (Gelens and Saurin, 2018). We therefore conclude that no efficient demethylase exists that affects global methylation levels in HeLa cells. Whether demethylation affects specific targets rather than the global ArgMet levels remains to be investigated.

We observed even in mouse tissues a large fraction of arginines being methylated with brain and spleen showing the highest ArgMet levels. In line with the specific pattern of ADMA, SDMA and MMA observed in cells, ADMA was the most abundant methylated species, followed by SDMA and MMA.

During ageing of mice, levels of protein ArgMet changed drastically in brain and spleen proteins, whereas other tissues, such as heart, liver and kidney, were less affected. The spleen is among the most affected organs during ageing, and a link to the accumulation of senescent cells has been hypothesised (Lim et al., 2010; Lim et al., 2008). Thus, it is conceivable that the loss of protein ArgMet is associated with loss of PRMT1 expression/activity under physiological conditions, as demonstrated by a recent study linking PRMT1 downregulation with senescence of neuroblastoma cells (Lee et al., 2019). High levels of protein ArgMet in spleen and a strong reduction during ageing raises the question whether accelerated ageing of the spleen could be an inevitable side effect of the aforementioned protein arginine methyltransferase inhibitors. First links between ArgMet and neurodegenerative diseases have been suggested as hypomethylated RNA-binding proteins FUS and poly-GR dipeptide repeats were found to be enriched in patients with frontotemporal dementia or amyotrophic lateral sclerosis, respectively (Dormann et al., 2012; Gittings et al., 2020; Suarez-Calvet et al., 2016). Given the globally reduced levels of SDMA/MMA in the brain of old mice, it will be interesting to investigate whether ArgMet is associated with risks of neurodegenerative diseases, for example through modulation of LLPS of RNA-binding proteins.

Taken together, our findings support the idea that (i) protein ArgMet is a highly abundant PTM in cells and tissues, (ii) ArgMet and specific aspects of metabolism are tightly coupled, (iii) demethylation is a slow process, and (iv) cancer and ageing lead to substantial changes in global ArgMet levels. Given its relatively high proportion, we hypothesise that ArgMet plays a key role in maintaining cellular homeostasis, e.g. by regulating LLPS and formation of membrane-less organelles on a global scale. Concentrations of ADMA, SDMA and MMA in proteins might be used as biomarkers for drug discovery, treatment response and (potentially) for diagnosis of tumour susceptibility for arginine methyltransferase inhibitors. These findings could lead to the development of novel and improved methods for basic research on ArgMet and implementing routine ArgMet-based screening in the clinic.

## Acknowledgements

The work was supported by Austrian Science Fund (FWF) grants P28854, I3792, and DK-MCD W1226 to T.M. and FWF grant P30162 to U.S.; T.M. was supported by the Austrian Research Promotion Agency (FFG) Grants 864690 and 870454; the Integrative Metabolism Research Center Graz; Austrian Infrastructure Program 2016/2017, the Styrian Government (Zukunftsfonds), and BioTechMed-Graz (Flagship project DYNIMO). D.K. was supported by the FWF (SFB F73, W1226, P32400, P30882, DP-iDP DOC 31), the BioTechMed-Graz flagship project “Lipases and Lipid Signaling”, the County of Styria, and the City of Graz. We thank the Center of Medical Research for access to cell culture facilities and Arno Absenger and Isabella Hindler for mouse care. F.Z. was trained within the frame of the PhD program Molecular Medicine, Medical University of Graz.

## Contributions

Conceptualization, T.M.;
methodology, F.Z., J.K.K., A.A., M.K., K.B.K., N.V., S.F., E.J-L., G.H., M.P., M.R.C, M.P., E.J.L., M.M;
software, F.Z., T.M.;
validation, F.Z., T.M.;
formal analysis, F.Z., T.M.;
investigation, F.Z., T.M.;
resources, T.M., G.H., E.S., D.K., U.S.;
data curation, F.Z., T.M.;
writing—original draft preparation, F.Z., T.M.;
writing—review and editing, F.Z., J.K.K., A.A., M.K., K.B.K., N.V., T.M., N.V., E.S., G.H. D.K., U.S., M.R.C., B.B., T.E., B.R., B.P., M.P., M.M.;
visualization, T.M., F.Z.;
supervision, T.M.;
project administration, T.M.;
funding acquisition, T.M., D.K, U.S‥

All authors have read and agreed to the published version of the manuscript.

## Methods

### Chemicals

L-arginine was obtained from AppliChem (Germany), ω-*N*^*G*^, *N*^*G*^-asymmetric dimethylarginine (ADMA), ω-*N*^*G*^-*N*’^*G*^-symmetric dimethylarginine (SDMA), and ω-*N*^*G*^-monomethylarginine (MMA) were obtained from Santa Cruz Biotechnology, Inc. (Germany). Sodium phosphate, dibasic (Na_2_HPO_4_), sodium hydroxide, chloroform, hydrochloric acid (32% m/v), and sodium azide (NaN_3_) were obtained from VWR International, 3-(trimethylsilyl) propionic acid-2,2,3,3-d_4_ sodium salt (TSP) from Alfa Aesar (Karlsruhe, Germany), deuterium oxide (^2^H_2_O) from Cambridge Isotope laboratories, Inc. (Tewksbury, MA). Adenosine, periodate oxidized (AdOx) and MS023 hydrochloride were obtained from Sigma Aldrich Austria. GSK591 (Synonyms: EPZ015866; GSK3203591 and GSK3368715 dihydrochloride (Synonyms: EPZ019997 dihydrochloride) were obtained from MedChemExpress Austria.

### Cell culture

MDA-MB-231 (Sigma Aldrich, Vienna, Austria), HaCat (ATCC, US), MCF10A (LGC Promochem, US), A375 (CLS, Germany), SW-872 (ATCC, US), 93T449 (ATCC, US), SW1353 (CLS, Germany) and juvenile fibroblasts fresh established from foreskin samples were obtained from the Center of Medical Research (ZMF), Medical University of Graz, Austria. HeLa (ATCC^®^, Guernsey, UK), fibroblast, HaCat, SW1353 and A375 cells were cultured in DMEM supplemented with 2 mM glutamine, 1% PS (100 U/mL penicillin, 100 μg/mL streptomycin) and 10% fetal bovine serum (FBS). MCF10A were cultured in DMEM with single quot kit suppl. Gr, 5% Horse Serum, 20 ng/mL hEGF, 0.5 μg/ml hydrocortison, 100 ng/ml choleratoxin, 10 μg/ml insulin and 2 mM glutamine. MDA-MB-231 were maintained in DMEM Hams F12 with 10% FBS, 2 mM glutamine and 1% PS. SW872 were cultured in DMEM Hams F12 supplemented with 5% FBS, 2 mM glutamine and 1% PS. 93T449 were cultured with RPMI-1640 with 10% FBS, 2 mM glutamine, 10mM 4-(2-hydroxyethyl)-1-piperazineethanesulfonic acid (HEPES), 1mM sodium pyruvate and 1% PS.

Cells were maintained in a humidified incubator at 37°C with 5% CO_2_. HeLa cells were treated for up to 3 days with AdOx (40 μM), MS023 (10 μM), GSK715 (2 μM), GSK591 (1 μM) and DMSO, before cell extracts were prepared.

### Yeast strain and culture conditions

Yeast experiments were carried out in *S. cerevisiae* BY4741 (*MATa his3Δ-1 leu2Δ-0 met15Δ-0 ura3Δ-0*) wild type yeast (Baker Brachmann et al., 1998) and the same strain carrying an *Hmt1*-knockout (Hmt1:kanMX), both obtained from Euroscarf. Cells were grown to logarithmic phase in SC 2% glucose medium consisting of 0.17% yeast nitrogen base (BD Difco™, 233520), 5% (NH_4_)_2_SO_4_ supplemented with 30 mg/L of all amino acids (except 80 mg/L histidine, 200 mg/L leucine and 26 mg/L methionine), 30 mg/L adenine, and 320 mg/L uracil. Fresh overnight cultures were diluted to 0.1 OD600 (Genesys 10uv photometer, corresponds to ~2×10^6^ cells/mL), incubated for 5 h at 28 °C, 145 rpm, shifted to medium lacking methionine and incubated for another 2 h to reach a culture density of ~0.6 OD600. 15 OD600 equivalents were harvested by centrifugation (1,700 g, 3 min, 4 °C), washed once with 10 ml ice-cold water, and the cell pellet was immediately snap frozen in liquid nitrogen and stored at −80 °C until processing for NMR analysis.

### Organoid culture

Mouse small intestinal organoids were cultured as described previously (Lindeboom et al., 2018). In short, the organoids were maintained using basic culture (ENR) medium, which contained advanced DMEM/F12 supplemented with penicillin/streptomycin (1%, 10 mM HEPES, 1× Glutamax, 1× B27 (all from Life Technologies) and 1 mM N-acetylcysteine (Sigma) supplemented with murine recombinant epidermal growth factor (Peprotech), R-spondin1-CM (5% v/v) and noggin-CM (10% v/v). A mycoplasma-free status was confirmed routinely. Organoids were split every 4–5 days by mechanical disruption and plated in Matrigel. Three days after splitting, stem cell‐enriched organoid cultures (CV) were generated by supplementation of ENR with CHIR99021 (3 μM) and valproic acid (1 mM). Paneth cell-enriched organoids were generated by addition of Chir (3 μM) and DAPT (5 μM), stem cell‐depleted organoid cultures (EN) were grown in ENR medium without R‐Spondin‐1. Organoids were harvested after 3 days by using mechanical dissociation of matrigel followed by 3 washing steps with ice‐cold PBS. Organoid pellets were immediately frozen at −80°C for further analysis.

### Animals and diets

For all experiments, young (9-11 weeks) and old (96-104 weeks) female wild type mice (mixed genetic background of 129/J and C57BL/6J) were used (n=5). Mice were maintained in a clean, temperature-controlled (22 ± 1°C) environment with a regular light–dark cycle (12 h/12 h) and unlimited access to chow diet (Altromin 1324, Altromin Spezialfutter GmbH, Lage, Germany) and water. All experiments were performed in accordance with the European Directive 2010/63/EU and approved by the Austrian Federal Ministry of Education, Science and Research (GZ 66.010/0051-WF/V/3b/2015).

### Sample preparation

Cells (5 x 10^6^) were plated onto 60 mm dishes and incubated under standard conditions as described above. To harvest the cells, medium was removed, cells were washed three times with 5 ml of cold phosphate-buffered saline (PBS, 137 mM NaCl, 2.7 mM KCl, 8 mM Na_2_HPO_4_, and 2 mM KH_2_PO_4_) solution and collected using a cell scraper. A solution of 5 × 10^6^ cells was centrifuged at 1,000 rpm for 1 min, the supernatant was discarded and the cell pellet was flash frozen in liquid nitrogen and stored at - 80°C for the extraction step. The organs were isolated from sacrificed mice, divided into 20–30 mg and snap-frozen in liquid nitrogen for storage at −80°C until extraction. Cell pellets, tissues and colon organoids were re-suspended in 400 μl ice-cold methanol (−20°C) and 200 μl MilliQ H_2_O and transferred to a tube containing Precellys beads (1.4 mm zirconium oxide beads, Bertin Technologies, Villeurbanne, France) for homogenization on a Precellys 24 homogeniser for 2 cycles of 20 seconds with 5,000 rpm, 10-s breaks. Cell and tissues debris were pelleted by centrifugation at 13,000 rpm for 30 min (4°C) and the precipitate was used for hydrolysis. Supernatants were frozen at −80°C and be used for e.g. metabolite analysis.

The precipitates were hydrolysed with 500 μl 9 M HCl for 12 h at 110°C to obtain (modified) amino acids. The solution was lyophilised and resuspended in 900 μl of 0.1 M HCl and 100 μl chloroform to remove lipids, centrifuged (10 min, 13,000 rpm) and the supernatant subjected to i) solid-phase-extraction (SPE) using Waters™ cartridges (1 ml Oasis MCX 1 cc/30 mg, Waters™, Eschborn, Germany) containing a mixed-mode polymeric sorbent with both reverse phase and cation exchange functionalities. Each step was performed with 1 ml of solution and by centrifugation at room temperature (1,000 rpm for 1 min). ii) auto SPE using Gilson^®^ GX-241 ASPEC system (Gilson Incorporated, Middleton, WI) and Waters™ cartridges. The flow rate for the injection of liquids was set to 2 ml/min for the sample, 7 ml/min for the replacement solution and the 0.1 M HCl, and to 10 ml/min for methanol, PBS and MilliQ-water. The cartridges were pre-conditioned with a detachment solution (2x 1 ml, 10% NH_3_ saturated solution, 40 % MilliQ H_2_O, 50 % methanol), methanol (1x 1 ml) and with PBS (2x 1 ml). After sample loading (1x 1 ml), cartridges were washed with MilliQ-water (3x 1 ml), 0.1 M HCl (5x 1 ml) and methanol (2x 1 ml,). The arginine and its derivatives were recovered with the replacement solution (2x 1 ml), lyophylized and dissolved in 500 μl NMR buffer [0.08 M Na_2_HPO_4_, 5 mM 3-(trimethylsilyl) propionic acid-2,2,3,3-d4 sodium salt (TSP), 0.04 (w/v) % NaN_3_ in D_2_O, pH adjusted to 7.4 with 8 M HCl and 5 M NaOH] for measuring.

### NMR measurements and spectral processing

All NMR experiments were acquired at 310 K using Bruker 600 MHz spectrometer equipped with a TXI probe head. The 1D CPMG (Carr–Purcell–Meiboom–Gill) pulse sequence (cpmgpr1d, 256 scans, size of fid 73728, 11904.76 Hz spectral width, recycle delay 4 s), with water signal suppression using presaturation, was recorded for ^1^H 1D NMR experiments. The 2D JRES (^1^H homo-nuclear J-resolved spectroscopy) pulse sequence (jresgpprqf, 64 scans, size of fid 16384/32, 10000.00/78.042 Hz spectral width in F2/F1, recycle delay 2 s) with presaturation during the relaxation delay was recorded to obtain virtually decoupled spectra. In brief, data were processed in Bruker Topspin version 4.0.2 using one-dimensional exponential window multiplication of the FID, Fourier transformation and phase correction. The spectral data were processed as previously described (Stryeck et al., 2018).

The ^1^H 1D projections of 2D J-resolved, virtually decoupled NMR spectra data processing was carried out using MestReNova 12.0.4 software’s automatic phase and baseline correction. Calibration was made by using tetramethylsilane (δH = 0). Quantification of arginine, MMA, ADMA and SDMA used integration of characteristic peaks. Calculation of absolute concentrations is based on known concentrations of external standards. The ADMA levels relative to total amounts of arginine are calculated by the formula: ADMA/arginine (%) = (integrals (ADMA)/integrals (arginine)) * (integrals (100μM arginine)/integrals (100μM ADMA)).

### Statistical analysis

Data are presented as mean ± standard deviation (SD) or standard error (SE). Statistical differences between control and AdOx treated groups (unpaired Student’s t test) or multiple groups (one-way ANOVA) are indicated by *P*-values of < 0.05 (*), < 0.01 (**), < 0.001 (***) or < 0.0001 (****). Statistical analyses and graphs were generated using Graph Pad Prism 5.01. software (GraphPad Software, La Jolla, CA, USA).

## Supplemental Information

**Figure S1.**
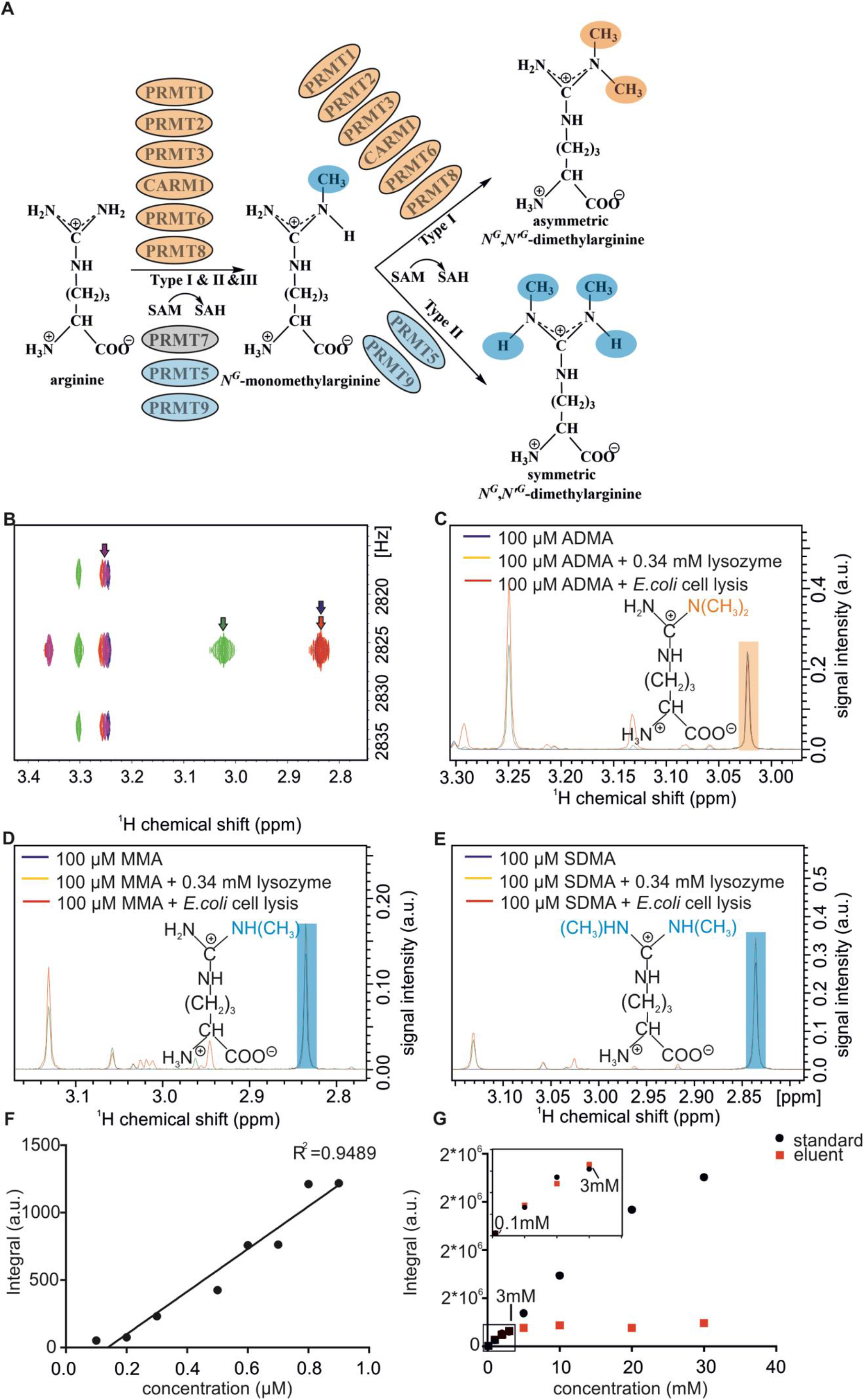
Related to Figure 1. Absolute quantification of protein arginine methylation by ArgMet-NMR. (A) Arginine residues are methylated by the protein arginine methyltransferase (PRMT) family members. PRMTs catalyze either the formation of monomethylarginine (MMA), asymmetric dimethylarginine (ADMA) or symmetric dimethylarginine (SDMA). (B) Overlay of ^1^H 2D J-resolved experiments of arginine (magenta), ADMA (green), MMA (purple) and SDMA (red), each 100 μM. (C) Overlay of ^1^H 1D projections of 2D J-resolved, virtually decoupled NMR spectra of ADMA recovery from lysozyme (yellow) and *E. coli* cell lysates (red). (D) Overlay of ^1^H 1D projections of 2D J-resolved, virtually decoupled NMR spectra of MMA recovery from lysozyme (yellow) and *E. coli* cell lysates (red). (E) Overlay of ^1^H 1D projections of 2D J-resolved, virtually decoupled NMR spectra of SDMA recovery from lysozyme (yellow) and *E. coli* cell lysates (red). (F) Correlation between integrals and concentration changes of ADMA signal of ^1^H 1D projections of 2D J-resolved spectra. Correlation coefficient (R^2^ = 0. 9489, p = 0.033) was computed with the Pearson Product Moment statistic. (G) Integrals of arginine recovery from SPE compared to standard concentrations.

**Figure S2.**
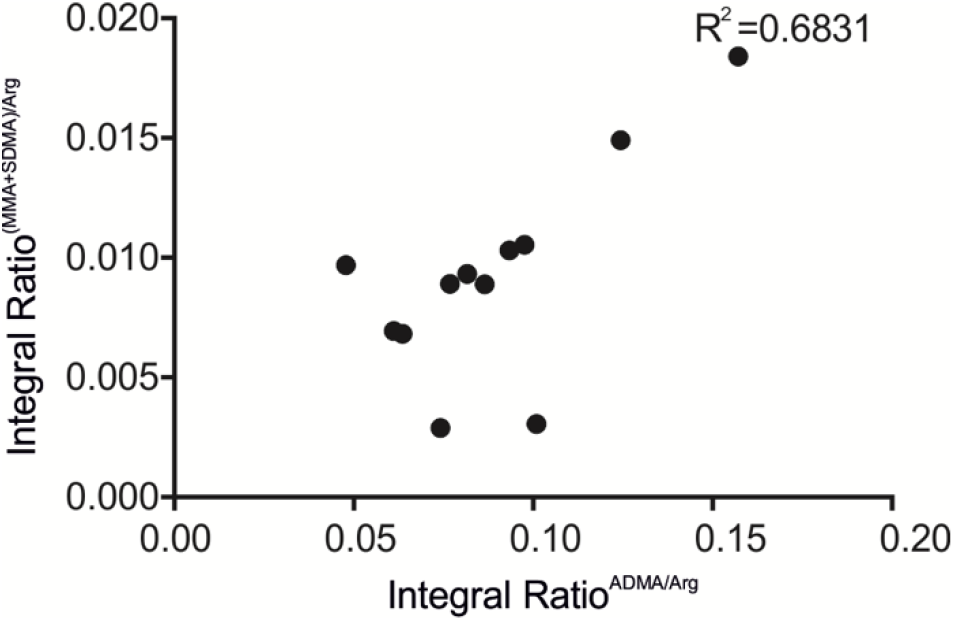
Related to Figure 2. Characterisation of ArgMet in purified proteins, yeast and mammalian cell lysates. The correlation between integral ratios of ADMA/arginine and (SDMA/MMA)/arginine. Correlation coefficient (R^2^ = 0. 6381, p = 0.0143) was computed with the Pearson Product Moment statistic.

**Figure S3.**
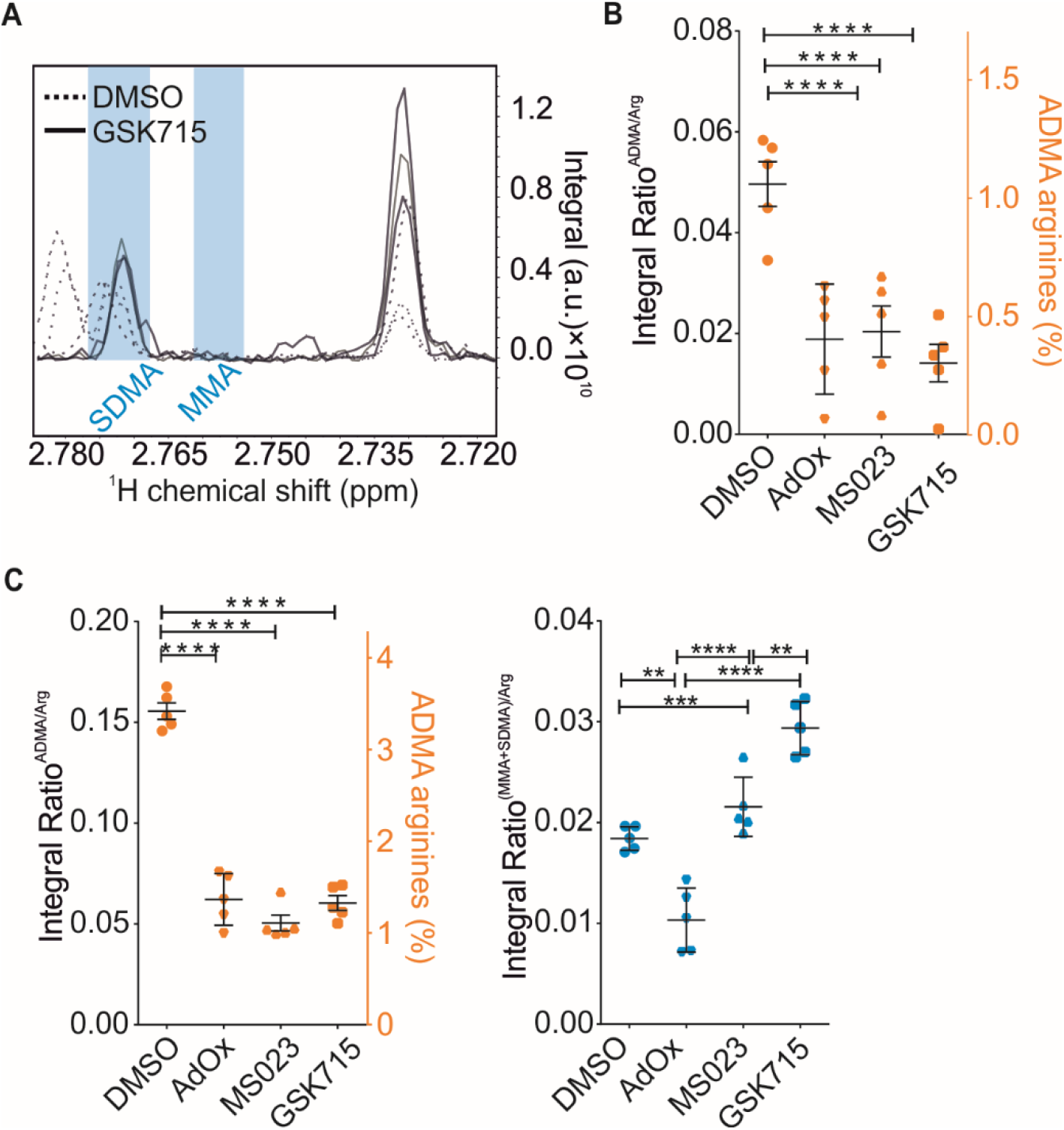
Related to Figure 3. ArgMet NMR enables quantification of protein ArgMet modulation and dynamics. (A) Spectral overlays of characteristic MMA and SDMA NMR methyl signals in d_6_-DMSO show that MMA and SDMA methyl resonances can be resolved (n=3), and higher SDMA represented in GSK715 than DMSO. (B) Protein ArgMet quantification of fibroblast cells treated for 3 days with either DMSO, 40 μM AdOx, 10 μM MS023, 2 μM GSK715 (n=5; mean ± SD). ADMA levels with respect to total amounts of arginine are shown. Both MMA and SDMA are non-detectable in these conditions. (C) Protein ArgMet quantification of A375 cells treated for 3 days with either DMSO, 40 μM AdOx, 10 μM MS023, 2 μM GSK715 (n=5; mean ± SD). ADMA levels are presented with respect to total amounts of arginine.

**Figure S4.**
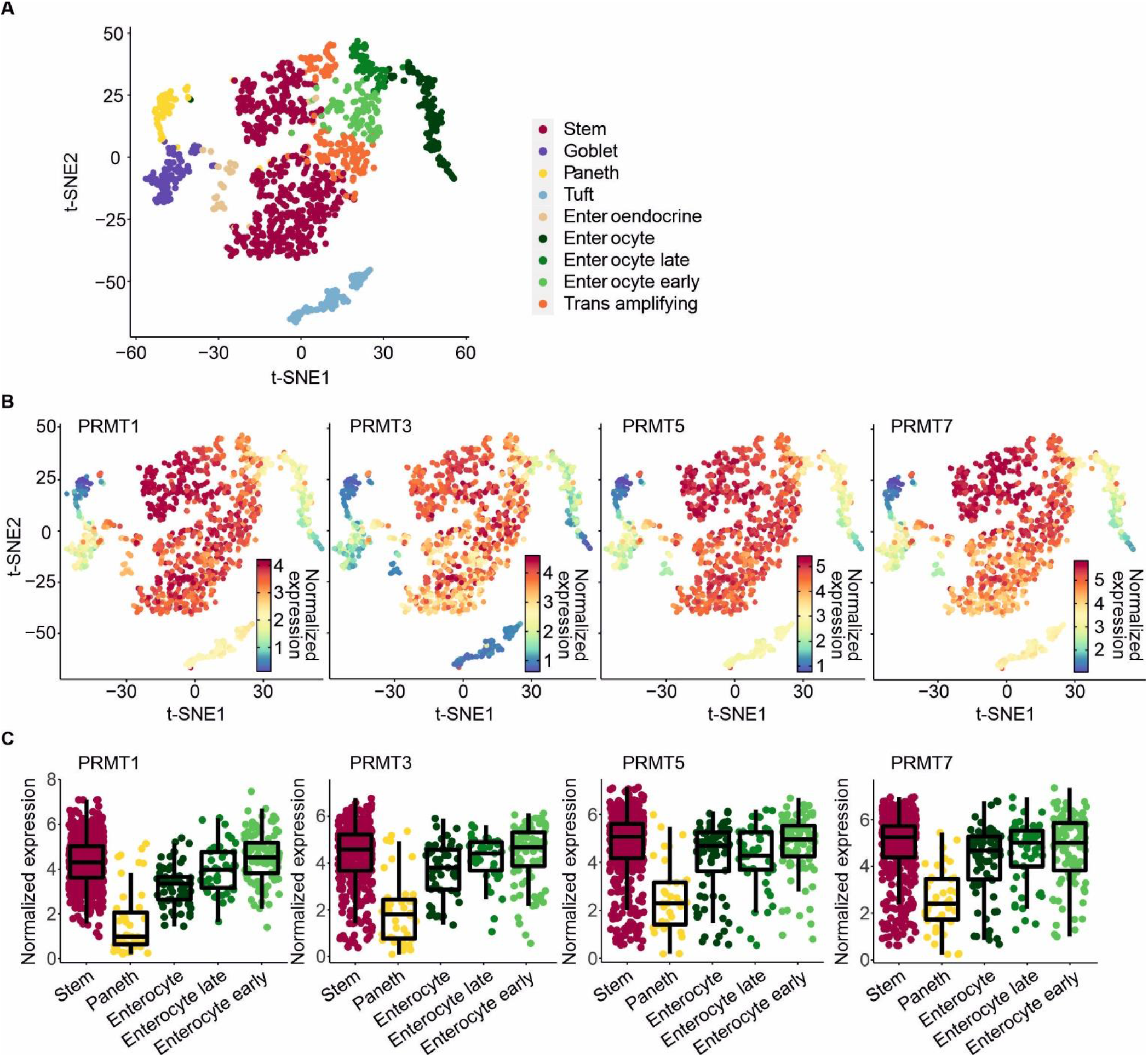
Related to Figure 4. NMR enables characterization of ArgMet in cell differentiation and ageing *in vivo*. (A) t-distributed stochastic neighbourhood embedding (t-SNE) visualization of 1,522 single-cell full-length sc-RNaseq data. Cluster annotation was done based on the expression of known cell type markers. (B) PRMT1, 3, 5 and 7 gene expression levels are plotted on t-SNE plots. (C) Boxplots represent the quantification of gene expression in each cluster gene (median values and 25th and 75th percentiles). The analysis was performed on mouse small intestine single cell RNAseq data set from Haber *et al.* (Haber et al., 2017) and the analysis was performed as described in Ludikhuize *et al.* (Ludikhuize et al., 2020).

